# Geometry and cell wall mechanics guide early pollen tube growth in *Arabidopsis thaliana*

**DOI:** 10.1101/2024.02.05.578915

**Authors:** Lucie Riglet, Catherine Quilliet, Christophe Godin, Karin John, Isabelle Fobis-Loisy

## Abstract

In *Arabidopsis thaliana*, successful fertilization relies on the precise guidance of the pollen tube tip as it navigates through the female pistil tissues to deliver non-motile sperm cells to ovules. While prior studies have unveiled the role of the pistil in directing pollen tubes to ovules, growth guidance mechanisms within the stigmatic epidermis during the initial phase of the pollen tube’s journey remains elusive. A recent analysis comparing wild-type (WT) pollen tube paths in WT and *ktn1-5* stigmatic cells revealed a tight connection between directed pollen tube growth and the mechanical properties of the invaded stigmatic cell. Building upon these observations, we constructed here a mathematical model to explore the mechanisms guiding early pollen tube growth through the papilla cell wall (CW). We found that in *ktn1-5*, the pollen tube moves freely on the curved papilla surface, following geodesics, while the WT papilla exerts directional guidance on the pollen tube. An order of magnitude analysis of the mechanical forces involved in pollen tube growth in papillae suggests a guidance mechanism, where the elongated papilla geometry and the CW elasticity combine to efficiently direct pollen tube growth towards the papilla base.

## I. INTRODUCTION

In the flowering plant *Arabidopsis thaliana*, reproduction initiates upon the arrival of a pollen grain at the receptive surface of the female organ, also called stigma (Fig. 1A,B). Once landed, the pollen grain germinates, forming a pollen tube responsible for transporting sperm cells to ovules deeply embedded within the pistil (Fig. 1C) [1]. Thus, precise guidance of the pollen tube is essential to ensure its correct path through the female tissues, preventing misrouting and securing the delivery of male gametes. While prior research has focused on pistil-produced chemical, electrical and mechanical signals directing pollen tubes towards the ovules [1–3], the early orientation of pollen tubes within the stigmatic epidermis has received limited attention. In *A. thaliana*, pollen tubes first penetrate the cell wall (CW) of the stigmatic cells (or papillae) and then travel, while being constrained within the rigid CW layer, towards the stigma base (Fig. 1D) [4–6].

**FIG. 1.**
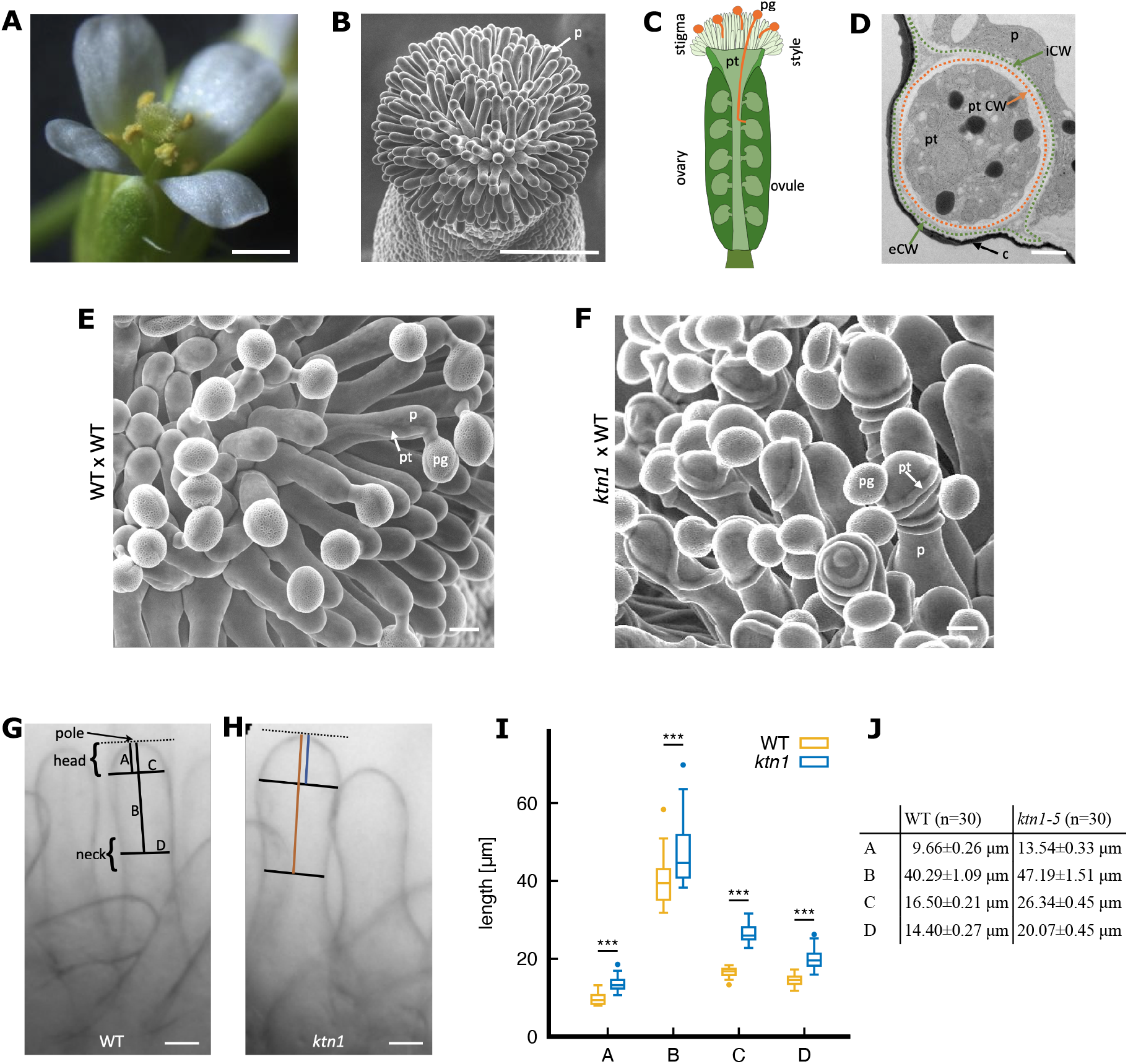
Pollen-stigma interaction in *A. thaliana* and characterization of papilla shape. (A) An Arabidopsis flower. The pistil at the center of the flower is surrounded by six anthers containing the male gametophytes (yellow pollen grains). Scale bar = 500 μm. (B) The pistil epidermis, the stigma, viewed by scanning electron microscopy (SEMi), is composed of dozens of elongated papillae (p). Scale bar = 100 μm. (C) Schematic representation of the pollen tube growth within the pistil. During pollination, pollen grains (pg) released from the anthers land on the stigma and germinate a pollen tube (pt). The tube transports the male gametes through the stigma, style and ovary towards the ovules for fertilization. (D) Transversal section of a pollinated papilla observed by transmission electron microscopy. The cuticle (c) appears as an electron dense black layer. The pollen tube progresses within the bilayered stigmatic CW inducing detachment of the internal (iCW) and external (eCW) layers. Stigmatic and pollen tube CW (ptCW) are highlighted with a green and orange dashed line respectively. Scale bar = 5 μm. (E, F) The pollen-stigma interaction views in SEMi. Most of the pollen tubes go straight towards the basis of the WT stigma (E) whereas pollen tubes make loops around the *ktn1-5* papillae (F). Scale bar = 10 μm. (G,H) Light microscopy images of WT (G) and *ktn1-5* papillae (H) with relevant shape descriptors A, B, C, and D. The head and neck regions are pointed out. Scale bar = 10 μm. (I) Box plots of dimensions A, B, C and D measured on 30 papillae from 3 WT or 6 *ktn1-5* stigmas. The horizontal bar in the boxes corresponds to the mean value. Statistical analysis was based on a non parametric Wilcoxon Rank Sum test. *** indicates a p-value<0.001. (J) Means and standard error of the mean (SEM) of the dimensions A, B, C and D. Measurements are provided in Table S2. n corresponds to the number of papillae analyzed. The label *ktn1* in (F, H, I and J) refers to the *ktn1-5* mutant.

Our recent work showed that the microtubule-severing enzyme KATANIN (KTN), by acting both on cortical microtubule (CMT) and cellulose microfibril (CMF) organization, conferred particular mechanical properties to the papilla CW, correlating with misguidance guidance of pollen tubes. On *ktn1-5* papillae, where the CW displayed impaired mechanical properties (i.e. isotropic CMT and CMF arrays, softer CW), wild-type (WT) pollen tubes exhibit a helical growth pattern (coiled path), occasionally growing in the opposite direction of the stigma base and ovules. In contrast, on WT papillae, characterized by anisotropic CMT and CMF arrays and stiffer CW, the WT pollen tubes maintain a relatively straight trajectory towards the stigma base (Fig. 1E,F). These correlations between the mechanical properties of the papillae CW and pollen tube paths suggest that mechanics could act as early guidance cue. Due to the technical challenges associated with a full characterization of the rheology and mechanical state of the papilla (wall stresses, turgor pressure…), we used phenomenological computational modeling to explore the mechanisms by which papilla may influence pollen tube growth. We tested our simulation by comparing the model predictions to experimental data from both our prior observations ([6]) and new experimental data concerning both WT and *ktn1-5* papillae. Our main results showed that the coiled paths of WT pollen tubes on *ktn1-5* papillae result from the absence of guidance and correspond to geodesic trajectories on the curved papilla surface. In contrast, the straight growth pattern observed on WT papillae requires a guidance cue that acts as a torque on the advancing pollen tube tip, redirecting its growth along the long papilla axis. We propose here a mechanism of mechanical growth guidance based on the elongated shape of the papilla and its CW elasticity. These minimal components are sufficient to explain how coiled and straight pollen tube growth arises and reproduce our experimental observations.

## II. RESULTS

### A. Shape differences in WT and *ktn1-5* papillae

Although both WT and *ktn1-5* papillae share an overall pin-like morphology with rotational symmetry, when looking closer, we noticed some shape differences, with WT cells apparently thinner than the *ktn1-5* ones (Fig. 1G,H). To investigate whether these potential disparities in papilla morphology play a role in impacting pollen tube growth trajectory, we first quantified WT and *ktn1-5* papilla shapes.

To that end, we defined geometric shape descriptors, as depicted in Fig. 1G,H. Papillae display a nearly spherical head region, characterized by measurements A and C (head length and width, respectively), which gradually transitions into a cylindrical shaft region that narrows at the neck. the papilla cylinder is characterized by the distances B (length from the papilla pole to the neck) and D (neck width).

We quantified these papilla shape descriptors and found significant differences (Fig. 1I,J and Suppl. Table SI). WT papilla cells are more slender with a narrower head (C=16.50 *μ*m ± 0:21 *μ*m) and neck (D=14.40 *μ*m ± 0:27*μ*m) than *ktn1-5* papillae (C=26.34 *μ*m ± 0:45 *μ*m; D=20.07 *μ*m ± 0:45 *μ*m), confirming our visual impression.

We then wondered whether these differences in papilla shapes could explain the difference in WT pollen tube trajectories, specifically the straight path within the CW of WT papillae *versus* the coiled paths within the *ktn1-5* papilla CW.

### B. Reference trajectories of a tube on a pin-like surface

We first investigated how variations in shape of a pin-like structure could influence the theoretical trajectories of an object moving along its surface. An object advancing on a curved surface without any external constraints follows the straightest line on this surface corresponding to the length-minimizing curve between pairs of points. By definition, this line is called a geodesic [7]. In other word, geodesics are to curved surfaces what straights lines are to planes. In addition, they possess the remarkable property that, at any specific point on the surface and in any given direction, there exists a single, unique geodesic.

We thus computed geodesic paths with fixed initial positions and directions on pin-like shapes with varying neck diameters, as illustrated in Fig. 2, by integrating differential equations describing geodesic trajectories on curved surfaces (details of the numerical procedure are given in **Methods**). Our simulations illustrate how the nature of the geodesic trajectories is sensitive to the actual surface shape of pin-like structures. As the neck diameter decreases, the geodesic changes. For instance, decreasing the neck diameter by 38% (from 0.43 to 0.27, see Fig. 2) resulted in markedly different coiling patterns (passing from 1 turn to 3.5 turns), especially around the narrowest region of the pin-like shape. In extreme scenarios, where the neck became exceedingly thin, geodesics could no longer cross the neck and remained confined to the upper part of the pin-like shape (rightmost two shapes in Fig. 2). Thus, according to this theoretical model, alterations in the pin-like shapes are sufficient to affect the path of an object moving along geodesics. However, it is important to notice that substantial alteration in shape are required to induce globally distinct trajectories.

**FIG. 2.**
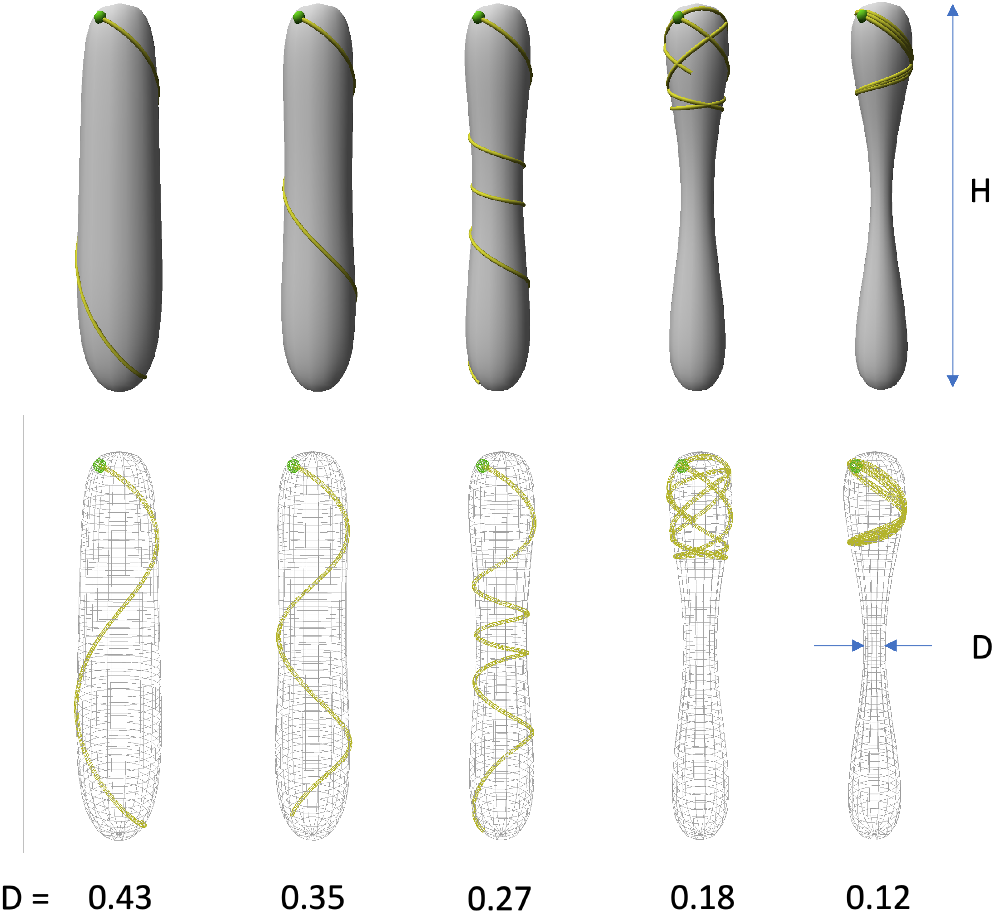
Geodesic trajectories on pin-like surfaces with varying shapes. From left to right: a series of pin-like shapes with decreasing neck mid-height diameter D and same height H. Top row: 3D surface rendering of the shape. Bottom row: wire-frame rendering where the whole geodesic trajectory is visible (yellow curve). All the geodesics start at the same position (green dot) at point *P*_0_ with azimuth = 0 and altitude = -0.1 with respect to the pole of the pin structure, and with an initial inclination angle of 28^°^downward with respect to the circumferential direction. D is given in arbitrary units (a.u.) and H=2.0 a.u. Note that the curve can cross itself (rightmost two situations).

### C. A minimal phenomenological mechanical model of pollen tube growth on the papilla surface

To further investigate the role of the papilla shape in pollen tube growth guidance, we developed a minimal phenomenological model integrating the experimentally measured geometries of both WT and *ktn1-5* papillae as well as known key features of the pollen tube growth.

Using the previously defined papilla descriptors (Fig. 1G,H), we fitted mathematical functions to derive analytical three-dimensional virtual WT and *ktn1-5* papillae surfaces **S** (Fig. 3A). Thereby papilla shapes are parameterized using cylindrical coordinates [*θ*; *z*], where *θ* and *z* represent the angular direction and the distance along the papilla axis (measured from the papilla pole), respectively.

**FIG. 3.**
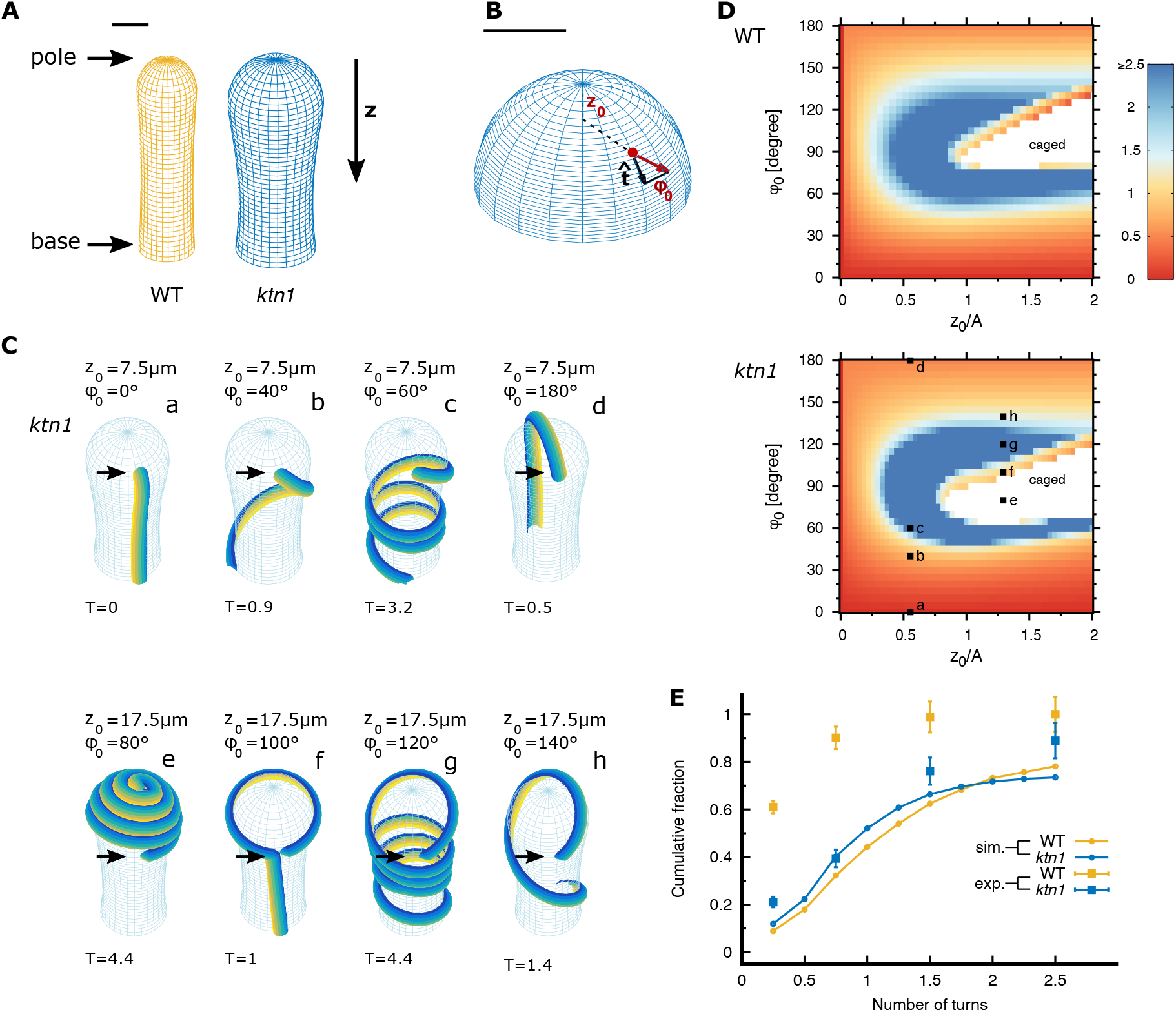
Papilla prototypes and pollen tube trajectories without guidance on WT and *ktn1-5* papillae. (A) Three dimensional shapes of WT and *ktn1-5* papillae. The papilla long axis is oriented along *z*. *z* = 0 corresponds to the papilla top. The scale bar is 10 *μ*m. (B) Three-dimensional view of the top region of a *ktn1-5* papilla with the initial position of the pollen grain (red dot) and initial direction of the pollen tube when emerged from the grain (red arrow), indicated as *z*_0_ and *ϕ*_0_ respectively. The vector 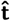 is a papilla surface tangent vector pointing in the longitudinal direction. 0^°^(180^°^) indicates an initial tube direction to the papilla base (tip). Scale bar = 10 *μ*m. (C) Examples of pollen tube trajectories on *ktn1-5* papilla surfaces, simulated with a combination of initial positions *z*_0_ (indicated by the black arrow) and directions φ_0_ of the pollen grain. Each configuration is represented by a letter from a to h. *T* stands for the number of turns the pollen tube makes to reach the papilla basis. Trajectories of pollen tubes on WT stigma with identical initial conditions are shown in Suppl. Fig. S2. (D) Morphological phase diagrams for the pollen tube turn number *T* depending on the initial pollen grain position *z*_0_ (normalized to the papilla distance A, see Fig. 1 G, H) and the initial pollen tube direction (φ_0_) on WT and *ktn1-5* papillae. *z*_0_*=A* = 0 denotes the papilla pole, *z*_0_*=A* = 1 the frontier between the head and cylindrical shaft and *z*_0_*=A* = 2 the pollen grain landing limit. The color code indicates the number of turns the trajectories undergoes before it reaches the papilla base. In the white region (caged) the tube path is caged by its own trajectory and cannot reach the papilla base; these trajectories were counted as trajectories with *T >* 2:5. The configurations depicted in (C) are indicated. (E) Comparison of simulated and experimental cumulative distribution of pollen tube turn number on WT (orange curve) and *ktn1-5* virtual papillae. Experiments were replotted from Riglet et al. (2020) [6], 192 pollinated papillae were analyzed; tube direction was classified in four categories (squares): straight, 0.5, 1 and 2 turns; the errorbars correspond to the SEM. The label *ktn1* refers to the *ktn1-5* mutant.

Given the peculiar geometry of the stigma, where papillae are fused at their base (Fig. 1B), and considering the substantial size of pollen grains released from the anthers (∼ 20 μm [8]), pollen grains land on the upper part of the papilla, above the neck. Moreover, numerous SEM observations have consistently shown the preferential attachment of pollen grains to the head region of the papilla (see as examples Fig. 1E,F and Suppl. Fig. S1). As a result, in our model, pollen grain attachment to the papilla and the corresponding landing position *z*_0_ is constrained to distances *z*_0_ *<* 2*A* (A distance shown on Fig. 1G), which corresponds approximately to the mid distance between the papilla pole and the neck. When emerges from the pollen grain, the pollen tube outgrowth has an initial direction, represented by the *φ*_0_ angle made with the longitudinal papilla axis. We assumed that this initial direction is devoid of any directional bias meaning that each possible initial direction are equally likely to occur. We adopted the convention that *φ*_0_ = 0 (*φ*_0_ = 180^°^) signifies that the tube grows along the long papilla axis towards the papilla base (pole), while an angle *φ*_0_ = 90^°^indicates an initial tube trajectory oriented along the circumferential direction of the papilla. The landing position *z*_0_ and the initial growth direction *φ*_0_ are schematically shown in Fig. 3B.

The pollen tube, a tip-growing cell, undergoes unidirectional elongation, with its expansion restricted to the very tip of the cellular protrusion [9]. Detailed mechanical models of tip growth consider the pollen tube as an elastic shell with an internal turgor pressure [10–12]. Here, in our phenomenological model, the pollen tube is considered as an inextensible filament with bending rigidity χ whose trajectory is confined to the papilla surface **S** by the CW bilayer (Fig. 1D and Ref. [13]). The pollen tube path is given by **X**(*s*) with *s* denoting the length of the trajectory from a given initial position. Denoting *L*(*t*) the length of the pollen tube at time *t*, the position of the elongating tip **P**(*t*) is given by the position of the filament extremity **P**(*t*) = **X**(*s* = *L*(*t*)). We assume that over a small time lapse δ*t*, the tip grows by a length δ*s* in a direction 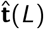, which results in a new tip position **P**(*t* + δ*t*) = **X**(*L*(*t* + δ*t*)) = **X**(*L*(*t*) + δ*s*).

To calculate the tip growth direction 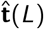, the tube bending 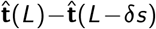 is decomposed into two components: (i) the bending normal to the surface to keep the tip trajectory on the papilla surface, and (ii) a lateral bending either to the left or to the right in the local tangent plane due to potential guidance forces (see Eq. (13) in **Methods**). Hence, in the absence of lateral forces, bending occurs only in the surface normal plane, and consequently the tube trajectory follows a geodesic.

An additional factor which influences the direction of tip growth is the self-avoidance property, i.e. an advancing tip cannot cross itself. This key feature of pollen tube growth, highlighted in our *in vivo* experiments on pollinated stigmas is illustrated in Suppl. Fig. S1. In our model, self-avoidance is implemented representing the pollen tube volume as a succession of spheres [at position **X**(*iδs*)] and cylinders [between positions **X**(*iδs*) and **X**(*iδs* + *δs*)] which cannot be penetrated by the outgrowing tip. In each simulation step, we tested that the growing tip does not penetrate into the excluded volume of a previously deposited pollen tube. When a potential penetration is detected, i.e. the advancing tip is in close proximity to a previously deposited tube path, the growth direction is not determined from the equation of momentum conservation (13) (see **Methods**). Instead we minimized the corresponding potential function (derived from (13) in **Methods**) under the constraint that the excluded volume is not violated.

### D. Pollen tube growth on *ktn1-5* papilla follows geodesics

Using the above computational model, we first simulated pollen trajectories on virtual papilla in the absence of lateral guidance cues from the stigmatic side. These geodesic trajectories are completely determined by the initial position, *z*_0_ and initial direction *φ*_0_ of the pollen tube tip. The behavior of pollen tube growth was assessed by calculating the number of turns T, as previously calculated in Ref. [6]. Experimentally, the turn number represents the number of revolutions made by a pollen tube around the papilla axis down to the observable papilla base (denoted as base in Fig. 3A), which corresponds to a distance of about *z* = 60 *μ*m.

We conducted simulations considering a broad range of initial pollen parameters *z*_0_ and *ϕ*_0_, and taking into account the self-avoidance property. Fig. 3C provides examples of simulated tube trajectories on *ktn1-5* papilla surfaces, along with the corresponding turn numbers *T*. For comparison, simulated trajectories on WT papillae are shown in Suppl. Fig. S3. Our simulations show that an initial growth direction *φ*_0_ close to the circular direction results in highly coiled paths (see Fig. 3C, (*z*_0_; *φ*_0_) = (7:5 *μ*m, 60^°^) and (*z*_0_; *φ*_0_) = (17:5 *μ*m, 120^°^)). However, this coiling can be mitigated if the tube tip is significantly deflected by its own path (see Fig. 3C, (*z*_0_; *φ*_0_ = (17:5 *μ*m, 100^°^)). Under specific initial conditions, the tube trajectory even self- constrains, as illustrated in Fig. 3C with (*z*_0_; *φ*_0_ = (17:5 *μ*m, 80^°^)), causing the tube to remain confined in the head region of the papilla.

To integrate the entire range of configurations we simulated, we employed a morphological phase diagram representation (Fig. 3D) illustrating the turn number depending on the initial pollen parameters (*z*_0_ normalized to the papilla distance A and *φ*_0_). Both morphological phase diagrams for WT and *ktn1-5* papillae display only minor disparities, suggesting that papilla shape differences (Fig. 1I,J) do not significantly impact the pollen tube trajectories. This is consistent with our earlier results for reference trajectories on a pin-like surface (Fig. 2), where we established that substantial shape differences are necessary to induce a change in the turning behavior of an object moving along geodesics.

Both phase diagrams show that trajectories starting from the papilla head region close to the pole with *z*_0_*=A* ≤ 0:25 consistently reach the bottom with relatively low turn numbers (orange color code in Fig. 3D). This mainly lies in the symmetry of the papilla head region; all initial angles *φ*_0_ lead to the same type of trajectory, corresponding to longitudinal lines, which either go directly to the papilla base (T ≈0) or cross over the papilla pole before going to the base (T ≈0:5). For trajectories starting at *z*_0_*=A* ≥0:25, the resulting tube paths highly depend on the initial tube direction, with a high turn number (blue color code) when a tube starts with an initial direction pointing in the circumferential direction (50^°^*< φ*_0_ *<* 130^°^). In this case trajectories enter the cylindrical papilla region at an oblique angle *‘* and then move either towards the papilla base in a helix (=geodesic on a cylindrical surface) or move towards the papilla pole, thereby confining themselves due to the self-avoidance behavior (see white regions in Fig. 3D). Trajectories starting with a low angle (φ *<* 30^°^) will move downwards with a turn number T ≈ 0 (see bottom regions in Fig. 3 D), whereas trajectories starting with a high angle (φ *>* 150^°^) will move over the papilla head region with a turn number T ≈ 0:5 (see top regions in Fig. 3D).

In the next step of our investigation, we conducted a quantitative comparison between simulated tube trajectories and experimental ones from Riglet et al. (2020) [6]. For a visually effective global comparison, we calculated the cumulative fraction of the number of turns T made by the pollen tube around the papilla for both our simulations and experimental datasets.

We found that cumulative fractions for simulated trajectories on both WT and *ktn1-5* virtual papillae (Fig. 3E, orange and blue solid curves) are very similar as expected from the phase diagrams (Fig. 3D). Around 40 − 50 % of the simulated pollen trajectories completed a single turn or less to reach the papilla base, whereas a smaller proportion of pollen tubes (around 20 %) reached the papilla base with a high turn number (more than 2.5). Simulated distributions closely resemble the experimental distribution of pollen tube turns on *ktn1-5* papillae (blue squares, Fig. 3E). In contrast, notable disparities were found when compared with the experimental trajectories pollen tubes make on WT papillae (orange squares, Fig. 3E).

In conclusion, our simulations of pollen tube growth in the absence of guidance cues indicate that (i) the difference in papilla shapes between WT and *ktn1-5* papillae are too small to account for different tube turning phenotypes and (ii) the parameters/conditions used in our simulations were sufficient to reproduce the highly coiling behavior of pollen tube trajectories experimentally observed on *ktn1-5* papillae. This suggests that pollen tubes on *ktn1-5* papillae grow following geodesics, without additional guiding cues from the stigma side.

### E. WT pollen tube trajectory deviation from geodesics requires guidance cues from the papilla side

We then wondered whether lateral forces deviating the pollen tube from geodesic paths could be at the origin of the low number of turns experimentally observed on WT papillae. We therefore introduced in our simulations an alignment force that orients the growth direction along the long papilla axis without any preferential orientation towards the papilla top or the base (nematic growth alignment) and which is proportional to sin 2φ, where *φ* denotes the angle between direction of the advancing tip and the papilla long axis. The magnitude of the lateral force is given by the adimensional constant μ, quantifying the ratio of the lateral forces and the internal resistance of the pollen tube against directional changes (provided by its bending rigidity). At this stage it is reasonable to assume that alignment forces exclusively operate in the cylindrical part of the papilla for *z* ≥ *A*, (Fig. 4A), since the papilla head region is likely isotropic in terms of its geometry and CW properties due to its nearly spherical shape.

**FIG. 4.**
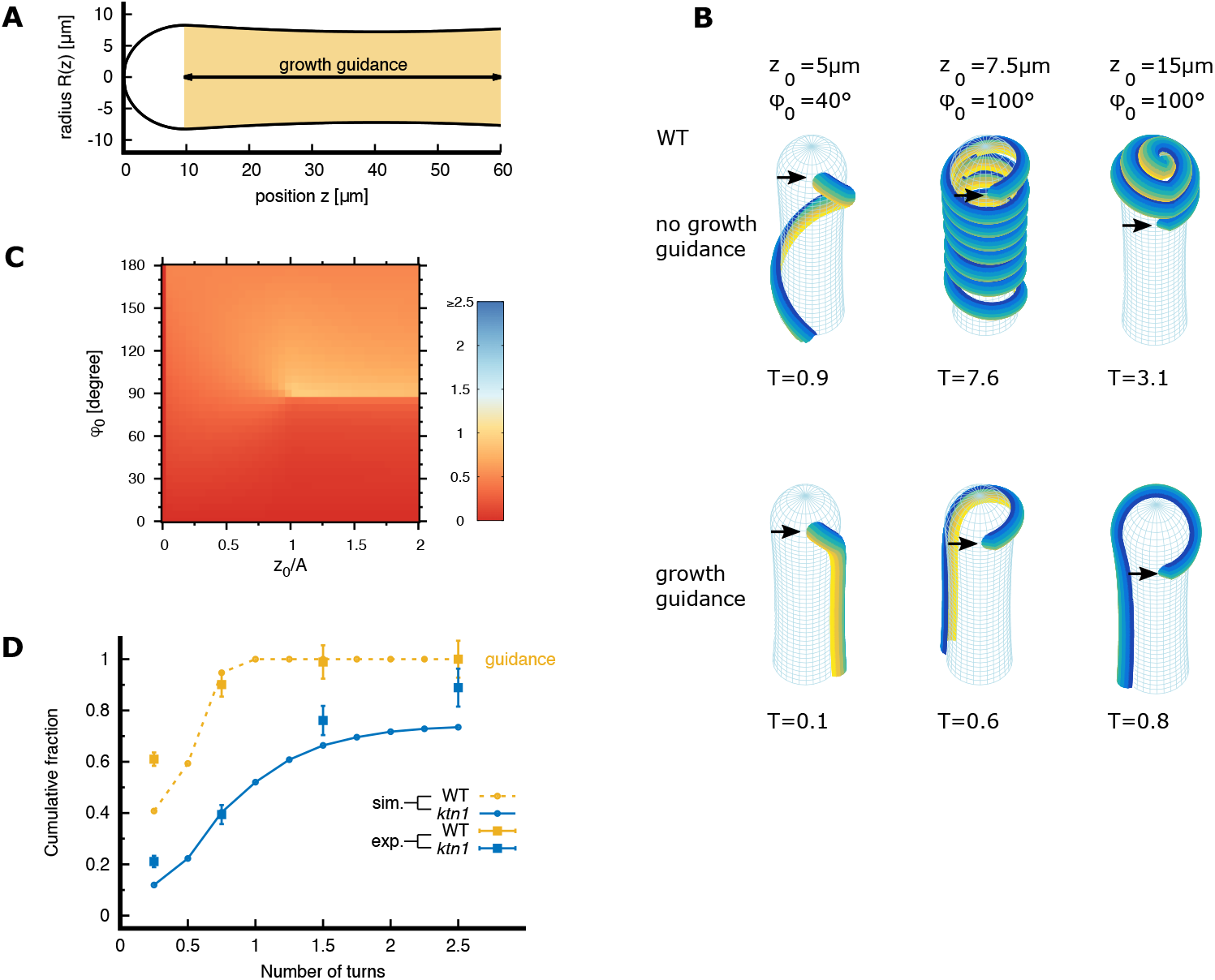
Effect of growth guidance (nematic alignment forces) on pollen tube trajectories. (A) Two-dimensional representation of the papilla, highlighting the papilla region where alignment forces act. (B) Example of pollen tube trajectories on WT papilla surface for various initial conditions *z*_0_ (indicated by the black arrow) and *φ*_0_, simulated without growth guidance (top) and with growth guidance (adimensional guidance strength *μ* = 0:1, bottom). T stands for the number of turns the pollen tube makes to reach the papilla basis. Trajectories of pollen tubes on *ktn1-5* papilla with identical initial conditions are shown in Suppl. Fig. S3A. (C) Morphological phase diagrams for the pollen tube turn number T depending on the initial pollen grain position *z*_0_ (normalized to the papilla distance A, see Fig. 1 G, H; *z*_0_*=A* = 0 denotes the papilla top most position) and the initial pollen tube direction (*φ*_0_) on WT papillae. The color code indicates the number of turns the trajectories undergoes before it reaches the papilla base. (D) Comparison of simulated and experimental cumulative distributions of pollen tube turn numbers. Simulated cumulative distributions for turn numbers on WT papillae are calculated with a growth guidance of *μ* = 0:1 (dashed orange curve). Simulated cumulative distributions for turn numbers on *ktn1-5* papillae are calculated without guidance (blue curve, reproduced from fig. 3E). Experiments were replotted from Riglet et al. (2020) [6], 192 pollinated papillae were analyzed; tube direction was classified in four categories (squares): straight, 0.5, 1 and 2 turns; the errorbars correspond to the SEM. The label *ktn1* refers to the *ktn1-5* mutant.

We performed numerical simulations of pollen tube trajectories using the same framework as in section II D (see also **Methods**) except that we incorporated the guidance factor *μ >* 0. The impact of alignment on pollen tube trajectories on WT surfaces is depicted in Fig. 4B. In configurations where pollen tubes were exhibiting significant coiling in the absence of growth guidance (Fig. 4B top row), they now follow straighter trajectories in the presence of a guiding cue (Fig. 4B bottom row), reaching the papilla base with minimal turning *T* ≤ 0:5. For instance, in the presence of guidance cues, tubes initially oriented towards the papilla base at a slight angle remain uncoiled (Fig. 4B bottom row, with *z*_0_ = 5 *μ*m and *φ*_0_ = 40^°^) and rapidly align with the long papilla axis. Trajectories initially oriented towards the papilla pole (Fig. 4B bottom row, with *z*_0_ = 15 *μ*m and *φ*_0_ = 100^°^) cross over the papilla head region and then reorient towards the papilla base. Notably, a guidance strength of *μ* = 0:1 along the long papilla axis is sufficient to direct pollen tubes towards the papilla base with a low number of turns (see phase diagram Fig. 4C; red-orange color for most of the initial conditions). For comparison, simulated trajectories and the phase diagram for *ktn1-5* papillae are shown in Suppl. Fig. S3 and confirm that shape differences between WT and *ktn1-5* papillae play only a minor role in pollen tube growth behavior, even in the presence of growth guidance from the papilla side.

When comparing the simulated and experimental cumulative distributions of turn numbers, a pollen tube growth model incorporating growth alignment with the long papilla axis of strength *μ* = 0:1 effectively reproduces the experimental turn number distributions observed on WT papillae (orange squares, Fig. 3E). For completeness, cumulative distributions for various alignment strengths μ are shown in Suppl. Fig. S3C.

In conclusion, we have here shown that a relatively weak guidance cue present in the papilla cylindrical region that aligns the pollen tube growth direction with the long papilla axis efficiently directs pollen tube growth towards the papilla base and reproduce the straight behavior of pollen tube trajectories experimentally observed on WT papillae.

### F. Estimation of the mechanical growth guidance through the papilla CW

We next questioned which mechanism could provide the guidance cue in the axial direction. As the pollen tube grows constrained between two CW leaflets, thus keeping it at the surface of the papilla without penetrating into the papilla volume, we reasoned that the guiding factor *μ* could originate from the papilla geometry itself, due to the local curvature anisotropy of the papilla surface.

Progression of the pollen tube provokes deformation of the papilla surface (inner/outer CW leaflets). Since the papilla is roughly a cylinder, this deformation depends on the tube growth direction (longitudinal *vs* circular). Indeed, when the pollen tube follows an axial direction, the CW is mainly strained in the circular direction, while the axial direction is not strained. By contrast, a pollen tube growing in the circular direction induces a CW deformation in both the circular and longitudinal directions (see Fig. 5A). Thus, pollen tube growth in the circular and the axial directions entails different energetic costs; axial growth being less costly than circular one. We hypothesized that this energetic difference that originates from the cylindrical form of the papilla could provide a strong guidance cue for a preferential tube growth in the axial direction, i.e. a higher elastic energy for growth in the circular direction compared to the axial direction.

**FIG. 5.**
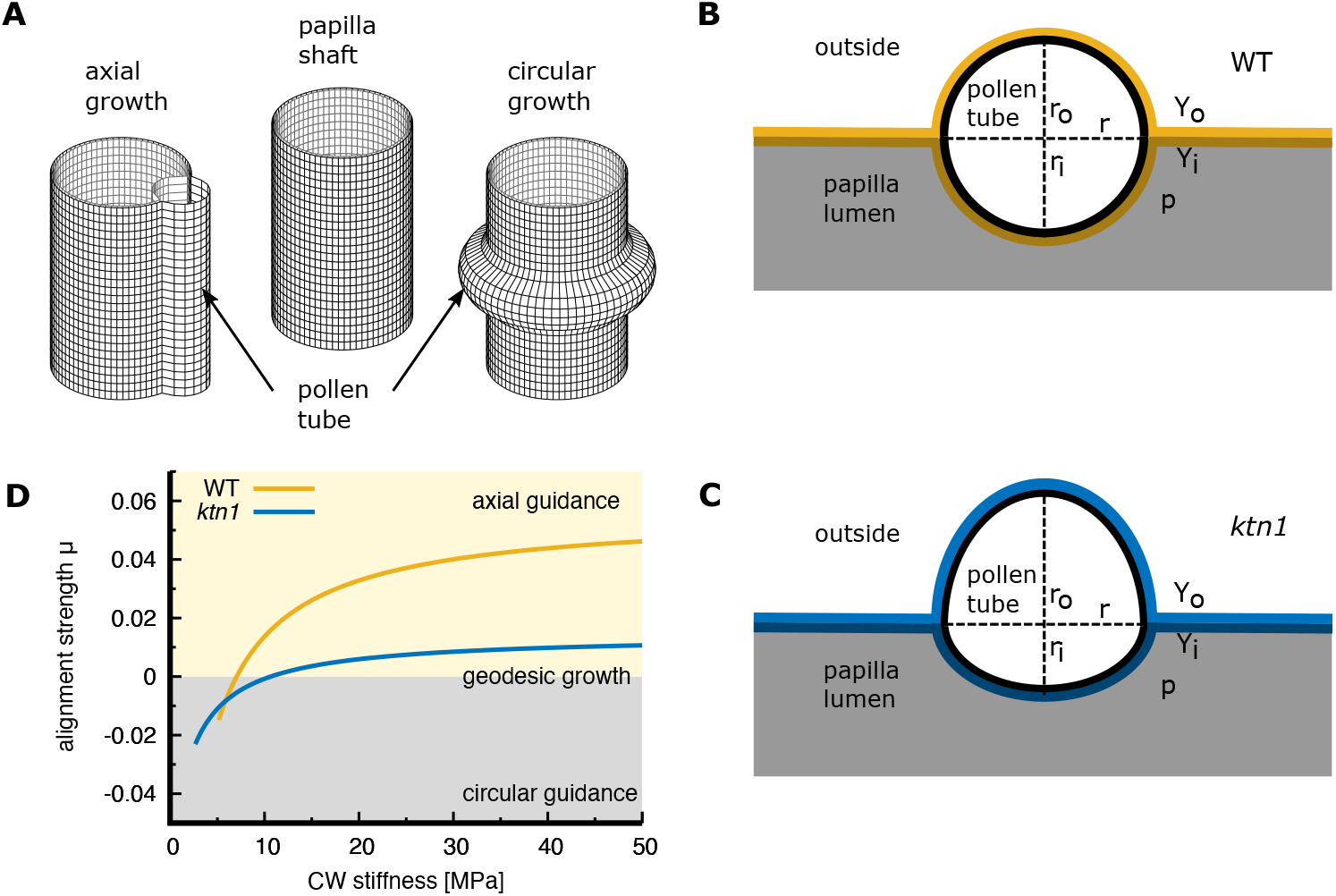
Model for pollen tube growth guidance by cell wall elasticity. (A) The orientation of pollen tube growth (axial *vs* circumferential) affects the strain in the papilla cell wall. (B,C) Mechanical model of pollen tube growth within the WT papilla CW (B) or the ktn1 papilla CW (C). A pollen tube (outer (inner) radius *r*_*o*_ (*r*_*i*_)) separates and deforms the CW bilayer with outer (inner) Young’s modulus *Y*_*o*_ (*Y*_*i*_) and performs volume work against the papilla pressure *p*. The shape of the pollen tube cross-section is approximated by two half-ellipses with the aspect ratio α = *r*_*o /*_ *r*_*i*_. (D) Adimensional alignment strength *—* for WT (α = *r*_*o /*_ *r*_*i*_ = 1) and *ktn1-5* (α = 3) papilla cells depending on the effective CW stiffness (*Y*_*i*_ + *Y*_*o*_)*=*2.

We further remarked that the tube growth generates a huge difference in CW deformations on WT and *ktn1-5* papillae [6]. In WT papillae, the lower half of the nearly circular pollen tube cross-section is immersed into the papilla volume (Fig. 5B and Ref. [6]), whereas in the *ktn* papilla, the indentation into the papilla volume is more shallow, corresponding to approximately one quarter of the pollen tube diameter (Fig. 5C and Ref. [6]). This suggests that the external CW leaflet of the mutant has a weaker rigidity relative to the inner wall. In contrast, the symmetrical indentation on the WT papilla suggests, that the outer CW is more rigid than the inner CW, since the stress in the outer CW has to work against the turgor pressure of the papilla. This conclusion is consistent with our previous AFM measurements, which showed that the WT CW is globally stiffer than that of the mutant *ktn1-5*.

We used our previous published data of the pollen tube indentation depths [6] to estimate the rigidity contrast between the two leaflets of the papilla CW (Suppl. Fig. S4). With these estimations we performed an order of magnitude calculation of the elastic energy for tube growth on the surface of a cylinder (internal pressure *p*) confined by two elastic layers (see **Supplementary Materials and Methods**). Thereby we considered two scenarios: growth in the circular with the elastic energy density *f*_*ci*_ and growth in the axial direction with the elastic energy density *f*_*ax*_ (see Fig. 5A). Within this picture, the adimensional alignment strength *μ* is given by *μ* ≈(*f*_*ci*_ − *f*_*ax*_)*l*^*2*^*/χ*, where *l* ∼ *r* denotes a typical length scale and χ denotes the bending rigidity of the pollen tube (**Supplementary Materials and Methods**).

Indeed we find that in a large range of CW rigidities, axial growth is more favorable than circumferential growth, thereby creating a torque on the advancing pollen tube tip that reorients pollen tube growth along the axial direction. Interestingly, the strength *μ* of this re-orientation cue is relatively independent of the CW stiffness as shown on Fig 5D), suggesting a robust behavior of the system working in the above ranges of parameters.

In addition, the model also provides an explanation for the different pollen tube behaviors in *ktn1-5* or WT CWs. Whereas a torque is present in both cases, the torque induced in the case of *ktn1-5* is markedly weaker (*μ* ≤ 0:01; Fig. 5D) than that induced in the WT (*μ* ≈ 0:05; Fig. 5D). As the alignment strength factor *μ* decreases, the turning behavior of the pollen tube increases (Suppl. Fig. S3C), and a *μ* factor of 0.01 may not be sufficient to reorient pollen tube towards the longitudinal axes.

Overall we can conclude that the cylindrical shape of the papilla and the CW elasticity combine to exert a torque on the advancing pollen tube tip, which acts as a guidance cue along the long papilla axis. The weak torque generated by the reduced CW rigidity in*ktn1-5* papillae, may not be sufficiently strong to deviate pollen tube growth from geodesics.

## III. DISCUSSION

The plant epidermis plays multiple essential roles and epidermal cells acquire a variety of shapes and sizes to perform specialized functions [13, 14]. The stigma covered with dozens of papillae serves as a receptive platform for pollen grains and the elongated shape of the stigmatic epidermal cells facilitates pollen capture for an efficient pollination process and reproduction success [15]. Here, we propose that the elongated cylindrical shape of the papilla (i.e. a high curvature anisotropy between the axial and circular directions) together with a tip growth constrained to keep at the papilla surface within the CW, is at the origin of a energetic difference for pollen tube growth that could provide an efficient guidance cue to favor axial growth and orient the pollen tube tip towards the papilla base.

The *ktn1-5* mutant, affected in CW properties, is known to exhibit alteration in plant cell shape [16, 17]. Quantifying WT and *ktn1-5* papilla shapes show that, while some differences exist, both papilla types share an overall cylindrical morphology. Despite this shared geometry, the comparison of simulated with experimental pollen tube growth trajectories revealed that the *ktn1-5* papillae provide less guidance cues for pollen tubes, which therefore make turns around the papilla, following geodesic lines. In addition, our simulation results excluded that shape differences are responsible for the observed differences in pollen tube growth behavior suggesting that, in addition to the geometry, an additional component would act to orient the pollen tube path and deviate substantially from geodesics in WT.

A unique feature of the pollen tube, notably when compared with other invasive organisms such as filamentous pathogens [18], is its ability to advance engulfed within the papilla CW throughout its journey towards the stigma basis. To progress within this rigid layer the pollen tube has to detach the inner and outer CW leaflets to form a cavity for growing, thus generating CW deformations.

An order of magnitude estimation of the mechanical energy associated with the papilla CW deformation during pollen tube growth indeed confirmed that the energetic costs for growth in the circular direction of the papilla is higher than axial growth. We showed that this difference in mechanical energy between the two growth directions leads to a torque which preferentially orients growth in the axial direction.

AFM measurements of CW elastic moduli and quantification of the CW deformation generated by the pollen tube growth [13] suggest that the *ktn1-5* mutant has a softer outer CW leaflet compared to the inner CW leaflet. Our order of magnitude estimations show that in this case, the mechanical guidance is too week to overcome the persistent pollen tube growth along geodesics.

In contrast, the WT exhibits a stiffer outer CW leaflet compared to the inner leaflet, which provides a sufficiently strong guidance cue to deviate pollen tube growth from geodesics and to efficiently orient the growth along the papilla axis. In conclusion, our simulations combined with quantitative comparisons with experimental data, reveal that a cylindrical papilla shape combined with sufficiently rigid CW leaflets, provide an efficient guidance cue along the axial papilla direction. This axial alignment is sufficient to prevent a coiled phenotype, akin to that observed in the *ktn1- 5* mutant. This suggests that a polar chemical gradient specifically guiding the pollen tube tip toward the base of the papilla cell is not necessary to reproduce the growth behavior observed on WT papilla cells.

## IV. METHODS

### A. Biological material and culture conditions

All *Arabidopsis thaliana* lines were in the Col-0 background and grown in growth chambers under long-day conditions (16 h light/8 h dark at 21^°^C/19^°^C with a relative humidity around 60 %). *ktn1-5* (SAIL_343_D12) was described previously [6]. All stigmas were analyzed at stage 13 of flower development [19].

### B. Measurement of shape parameters of papilla cells

Stigma were observed with an upright optical microscope Zeiss Axioimager Z1, under bright field, using a Plan-Apochromat 10x objective. Images processing and dimension measurements were performed with the ImageJ/Fiji software [20].

### C. Differential equations describing geodesic lines on a parametric surface

#### a. Geometric model of a pin-like surface

In Fig. 2, we represented pin-like shapes as surfaces of revolution. These surfaces were formalized by defining a profile curve represented as a NURBS curve *N*(*u*^1^) [21], *u*^1^ being a free real parameter between 0 and 1, that is then rotated around a vertical axis according to the equation

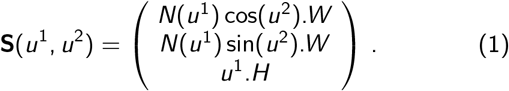

**S**(*u*^1^; *u*^2^) represents a point at the surface, corresponding to parameters (*u*^1^; *u*^2^). Here, *u*^1^ represents a normalized *z* coordinate - between 0 and 1 - pointing upward and *u*^2^ represents the azimuthal coordinate varying in [0; 2π[. The NURBS curve *N*(*u*^1^) is defined using a set of control points and scales in width and height, respectively denoted by scalars *W* and *H*. Different surface profiles with varying radii at mid-height (referred to as ‘the neck’) of the model were obtained by changing the most central control point CP3 (see Suppl. Table SII).

#### b. Geodesics

Let us consider a parametrized surface **S** and a coordinate system (*u*^1^; *u*^2^) on this surface. The points *P* of the surface in ℝ^3^ are defined by their coordinates *x*^*i*^ (*u*^1^; *u*^2^), *i* = 1; 2; 3.

A curve embedded in the surface is defined as a continuous mapping γ : *t* → *x*^*i*^ (*u*^1^(*t*); *u*^2^(*t*)). A geodesic on the surface is a curve whose length is locally minimal between any two of its points. It can be shown that geodesics are also the straightest curves that can be drawn on the surface (lines with least total curvature), e.g. see [22].

On a smooth surface, geodesics have the following remarkable property: given a point *P* = [*u*^1^; *u*^2^]^*T*^ on the surface and a vector **V** in the tangent plane at *P*, there exists a unique geodesic curve starting at *P* whose tangent is in the direction **V**, [22].

The curve γ is arc-length parametrized (parameter *s*) using a smooth mapping γ(*s*) = *x*^*i*^ (*u*^*α*^(*s*)) from a real interval 𝕀 on the curved space 𝒮, such that, as *s* varies, the point *γ*(*s*) travels at a constant and unit velocity:

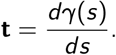

At a given point *P*, the variation of this tangent vector on the surface with *s* defines the curvature vector:

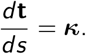

This vector is not necessarily in the tangent plane at *P*. In general it can be decomposed into normal and in-plane components at *P* :

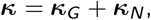

where **κ**_*G*_ is the in-plane component vector of the curvature, called the geodesic curvature, and **κ**_*N*_ is the normal curvature vector (aligned along the normal to the surface at *P*). A curve such that the geodesic curvature is null at every point is called *a geodesic*, meaning that the curve does not bend in the local tangent plane at any point, while it may bend in the direction normal to the surface (**κ**_*N*_ *≠* **0**), [22].

This definition can be used to derive the equation of geodesic curves on 𝒮 which is given by a set of two second order, non-linear, coupled differential equations, one for each value of α, e.g. [22]:

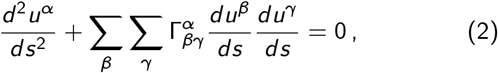

where 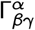 are the Christoffel symbols of the second kind computed from the first and second derivatives of the surface equation and from scalar products between these. Using Eqs. 2, it is possible to compute geodesic trajectories on the surface from given initial conditions corresponding to some initial position and orientation on the surface. This defines an initial value problem. A classical strategy to solve such an initial value problem with second order differential equations similar to Eqs. 2, consists of considering 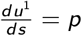 and 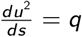 as two new independent variables and rewrite Eqs. 2 as a system of four coupled first order differential equations [23]:

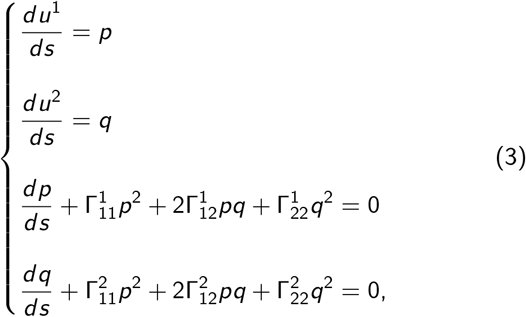

with the initial conditions:

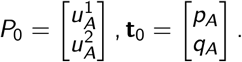

In Fig. 2 we used *P*_0_ = [0; − 0:1]^*T*^ (the point is slightly below the tip) and an initial inclination is **t**_0_ = [6:7; − 0:47]^*T*^ corresponding to an angle of 28^°^downwards with respect to the horizontal. To integrate these equations we used the odeint function of the Python SciPy library [24].

### D. Analytical expressions for virtual papilla shapes

To proceed further with the geometric model of the papilla surface and relate its parameters to geometric quantities that can be measured on the images, we designed a variant of the previous parametric expression of the surface, where the dependency of the papilla radius *R* on the z coordinate is expressed by an explicit formula with easily interpretable geometric parameters. As in the previous section, we expressed the rotational symmetry of the papilla surface **S** using a surface of revolution with cylindrical coordinates (θ, *z*) as:

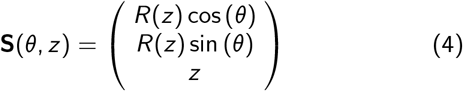

with the papilla long axis oriented along *z*. However, the radius function *R*(*z*) is now given by:

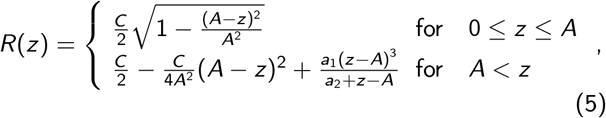

where the parameters *A* and *C* can readily be measured on the images, (cf. Fig. 1G,H). Note that the papilla head is described as an ellipsoid. At *z* = *A* the radius function is continuous up to the second derivative of *R* w.r.t. *z*. The shape parameters *a*_1_ and *a*_2_ were adapted to fulfill the following conditions (cf. Fig. 1G,H)

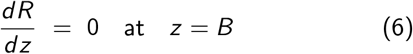

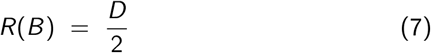

and can be calculated from the following analytical expressions

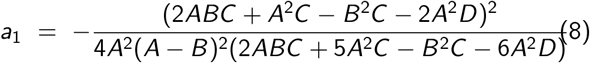

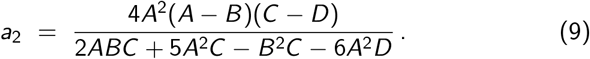

### E. Numerical details of the phenomenological model for pollen tube trajectories

The pollen tube path **X**(*s*) is discretized into equidistant points **X**(*iδs*), where *s* denotes the arc length.

#### a. Cylindrical papilla part

In the cylindrical region of the papilla [i.e. where the surface can be conveniently parametrized in cylindrical coordinates (θ, *z*), see Eq. (4)] the pollen tube path is given by the following ordinary differential equation

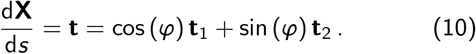

*φ* denotes the angle between the tip tangent **t** and the surface tangent 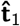 pointing along the long papilla axis

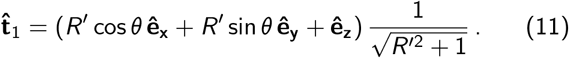

In the here chosen definition of *φ*, an angle *φ* = 0^°^(*φ* = 180^°^) indicates a pollen tube tip oriented towards the papilla base (pole).

The second surface tangent 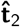 is oriented in the circumferential direction of the papilla surface and forms an orthonormal basis with 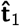

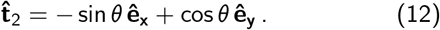

The momentum conservation at the pollen tube tip is given by the following ordinary differential equation and determines the evolution of the angle *φ*(*s*) along the pollen tube trajectory

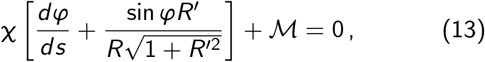

where χ denotes a bending rigidity and where

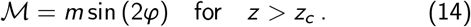

denotes the external torque of magnitude *m* which aligns the growth direction (described by *φ*) with the long papilla axis for *z > z*_*c*_. Here we chose *z*_*c*_ = *A*, where the almost spherical cap of the papilla goes over to the cylindrical shaft. Note that expression (14) favors growth along the axial direction. Depending on the angle *φ* the preferred growth direction is towards the papilla pole or the papilla base. *R*^′^ denotes the first derivative of *R* w.r.t. *z*, i.e. 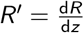. Eqs. (13) can be derived from a variational principal considering the following energy functional for the tube bending energy

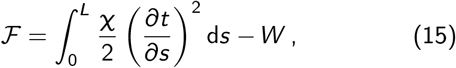

where *W* denotes the work performed by external forces on the tube tip. Variation of ℱ w.r.t. the tube tangent vector **t** results in

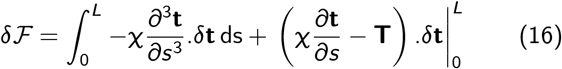

The bulk term and the boundary term at *s* = 0 in Eq. 16 vanishes since the tube cannot change its position after it has been deposited. With **T** = 2*m*(**t**:**t**_1_)**t**_1_ and using cylindrical coordinates we recover Eqs. (13) and (14).

To express the alignment strength *m* compared to the tube bending rigidity χ we introduce the adimensional growth guidance *μ* = *m𝓁 / χ*, with *𝓁* being the typical length scale (here *𝓁* ∼ 2:5 *μ*m, corresponding to the pollen tube radius). In the absence of any external torque, i.e. ℳ = 0, Eqs. (10) and (13) describe a geodesic line on the papilla surface **S**, i.e. the component of the curvature vector of the tube trajectory 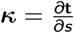 that lies in the tangent plane of the surface **S** is null.

*b. Papilla head region* In the head region of the papilla, i.e. *z* ≪ *A* the parametrization of the tube path in cylindrical coordinates will fail. Here we have parametrized the papilla surface in Cartesian coordinates (*x; y*)

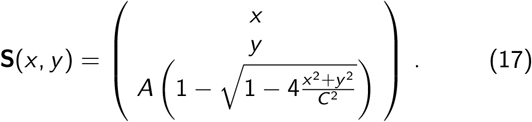

Then a choice of orthonormal surface tangent vectors is

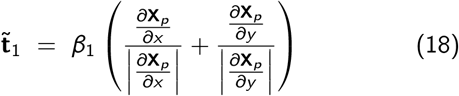

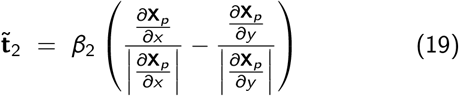

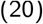

where *β*_1_ and *β*_1_ denote normalization factors. Since guidance cues are absent in the cap region the momentum conservation reduces to

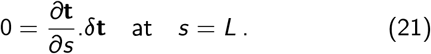

For simplicity, in the head region we have used the discrete form of (21)

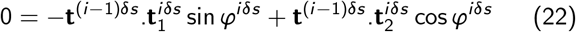

to determine the new tangent direction *φ*^*iδs*^ at the position of the tube extremity **X**(*iδs*).

#### c. Implementation of self-avoidance

While Eq. (13) can be numerically integrated in the absence of any self-avoidance, it is less practical to handle if self-avoidance plays a role. Instead, in situations where the growing tip was likely to cross over its previously deposited tube, we calculated the minimum of an effective potential ℱ_*tip*_ with the constraint, that the tip of the tube is not penetrating an existing tube path with radius *r* = 2:4 *μ*m [25]. The tip potential ℱ_*tip*_ in its discrete form is given by

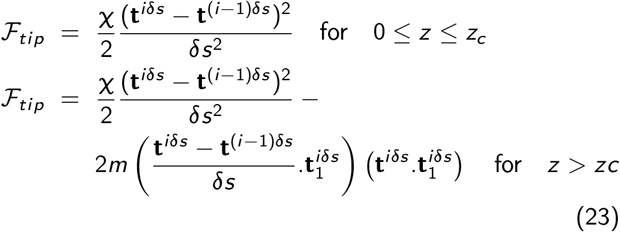

Minimizing the potential ℱ (23) w.r.t. to the angle *φ*^*s*^ by respecting self-avoidance then determines the tip direction. In a small region near the papilla pole the unit surface tangents 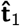 and 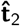 have to be replaced by the unit surface tangents 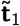 and 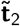. The pollen tube path itself is described as a succession of spheres [at position **X**(*iδs*)] and cylinders [between positions **X**(*iδs*) and **X**(*iδs* + *δs*)] which represent an excluded volume, which cannot be penetrated by the outgrowing tip. In each simulation step, we tested that the growing tip does not penetrate into the excluded volume of a previously deposited pollen tube. When a penetration was possible, the growth direction was determined from minimizing a potential function (23) under the condition that the excluded volume is not violated.

### F. Calculation of cumulative distributions of turn numbers

To calculate the cumulative distributions of turn numbers *T* from morphological phase diagrams (see Figs. 3 and 4) we assumed that pollen grains can only attach to the papilla cell in the upper part of the papilla for 0 ≤ *z*_0_ ≤ 2*A*. Furthermore, the turn number of each initial attachment site *z*_0_ is weighted in the cumulative distribution with a factor 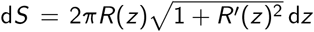 corresponding to the *z* - dependent surface area element of the papilla. Trajectories which could not reach the papilla base due to self-avoidance were always counted as trajectories with high turn numbers (T*>* 2:5). Typically, morphological phase diagrams where calculated for increments in the starting angle *Δ*φ_0_ = 5^°^and the starting position Δ*z*_0_ = 5 *μ*m.

## Notes

### Competing Interest Statement

The authors have declared no competing interest.

## References

[1] Fobis-Loisy, I. & Jaillais, Y. Feeling the pressure: a mechanical tale of the pollen tube journey through the pistil. Developmental Cell 56, 873–875 (2021).

[2] Mizuta, Y. & Higashiyama, T. Chemical signaling for pollen tube guidance at a glance. Journal of Cell Science 131, jcs208447 (2018).

[3] Clarke, D., Morley, E. & Robert, D. The bee, the flower, and the electric field: electric ecology and aerial electroreception. Journal of Comparative Physiology A 203, 737–748 (2017).

[4] Kandasamy, M. K., Nasrallah, J. B. & Nasrallah, M. E. Pollen-pistil interactions and developmental regulation of pollen tube growth in arabidopsis. Development 120, 3405–3418 (1994).

[5] Elleman, C., Franklin-Tong, V. & Dickinson, H. Pollination in species with dry stigmas: the nature of the early stigmatic response and the pathway taken by pollen tubes. New Phytologist 121, 413–424 (1992).

[6] Riglet, L. et al. Katanin-dependent mechanical properties of the stigmatic cell wall mediate the pollen tube path in arabidopsis. Elife 9, e57282 (2020).

[7] Bronstein, I. N., Semendjajew, K. A., Musiol, G. & Muehlig, H. Handbook of Mathematics (Springer, 2015).

[8] Rozier, F. et al. Live-cell imaging of early events following pollen perception in self-incompatible arabidopsis thaliana. Journal of Experimental Botany 71, 2513–2526 (2020).

[9] Dumais, J. Mechanics and hydraulics of pollen tube growth. New Phytologist 232, 1549–1565 (2021).

[10] Fayant, P. et al. Finite element model of polar growth in pollen tubes. The Plant Cell 22, 2579–2593 (2010).

[11] Kroeger, J. H. & Geitmann, A. Pollen tube growth: Getting a grip on cell biology through modeling. Mechanics Research Communications 42, 32–39 (2012).

[12] Bidhendi, A. J. & Geitmann, A. Finite element modeling of shape changes in plant cells. Plant Physiology 176, 41–56 (2018).

[13] Riglet, L., Rozier, F., Fobis-Loisy, I. & Gaude, T. Katanin and cortical microtubule organization have a pivotal role in early pollen tube guidance. Plant Signaling & Behavior 16, 1921992 (2021).

[14] Riglet, L., Gatti, S. & Moyroud, E. Sculpting the surface: Structural patterning of plant epidermis. iScience 24, 103346 (2021).

[15] Dresselhaus, T. & Franklin-Tong, N. Male–female crosstalk during pollen germination, tube growth and guidance, and double fertilization. Molecular Plant 6, 1018–1036 (2013).

[16] Bichet, A., Desnos, T., Turner, S., Grandjean, O. & Hofte, Botero1 is required for normal orientation of cortical microtubules and anisotropic cell expansion in arabidopsis. The Plant Journal 25, 137–148 (2001).

[17] Burk, D. H., Liu, B., Zhong, R., Morrison, W. H. & Ye, Z.-H. A katanin-like protein regulates normal cell wall bio-synthesis and cell elongation. The Plant Cell 13, 807 (2001).

[18] Riglet, L. et al. Invasion of the stigma by the pollen tube or an oomycete pathogen: striking similarities and differences. bioRxiv (2023).

[19] Smyth, D. R., Bowman, J. L. & Meyerowitz, E. M. Early flower development in arabidopsis. The Plant Cell 2, 755–767 (1990).

[20] Schindelin, J. et al. Fiji: an open-source platform for biological-image analysis. Nature methods 9, 676–682 (2012).

[21] Piegl, L. & Tiller, W. The NURBS Book (CRC Press, 1997).

[22] Kreyszig, E. Differential geometry (Dover, 1991).

[23] Gray, A. Modern Differential Geometry of Curves and Surfaces with Mathematica, Second Edition (CRC Press, 1997).

[24] Virtanen, P. et al. SciPy 1.0: Fundamental Algorithms for Scientific Computing in Python. Nature Methods 17, 261–272 (2020).

[25] Riglet, L. et al. Invasion of the stigma by the pollen tube or an oomycete pathogen: striking similarities and differences. bioRxiv 2023.07.19.549726 (2023).

